# Hoxb5 reprograms murine multipotent blood progenitors into hematopoietic stem cell-like cells

**DOI:** 10.1101/2021.10.25.465815

**Authors:** Dehao Huang, Qianhao Zhao, Qitong Weng, Qi Zhang, Kaitao Wang, Lijuan Liu, Chengxiang Xia, Tongjie Wang, Jiapin Xiong, Xiaofei Liu, Yuxian Guan, Yang Geng, Fang Dong, Hui Cheng, Jinyong Wang, Mengyun Zhang, Fangxiao Hu

## Abstract

The expression of transcription factor Hoxb5 specifically marks the functional hematopoietic stem cells (HSC) in mice. However, our recent work demonstrated that ectopic expression of Hoxb5 exerted little effect on HSC but could convert B cell progenitors into functional T cells in vivo. Thus, cell type- and development stage-specific roles of Hoxb5 in hematopoietic hierarchy await more extensive exploration. Here, with a mouse strain engineered with conditional expression of Hoxb5, we unveiled that induced expression of Hoxb5 in mouse multipotent progenitor cells (MPP) led to the generation of a *de novo* Sca1^+^cKit^+^Mac1^+^CD48^+^ (Mac1^+^CD48^+^SK) cell type, which has the ability to repopulate long-term multi-lineage hematopoiesis in serial transplant recipients. RNA-seq analyses showed that Mac1^+^CD48^+^SK cells exhibited an acquired machinery of DNA replication and cell division, which resembled nature fetal liver HSC cells (FL HSC). In short, our current study uncovers that Hoxb5 is able to empower MPP with self-renewal potential, thereby providing new strategies to reprogram blood progenitor cells into HSC-like cells.

## Introduction

Hematopoietic stem cell (HSC) is the blood cell type that possesses dual features of self-renewal and multi-lineage potential, which are critical for replenishing the entire hematopoietic system throughout an individual lifespan (Morrison and Weissman, 1994; Seita and Weissman, 2010). However, the absolute numbers of HSC in adults are extremely rare (Abkowitz et al., 2002; Bernitz et al., 2016) and are not efficiently expanded in vitro (Kumar and Geiger, 2017; Tajer et al., 2019). Researchers have been attempting alternative approaches to generate engraftable blood progenitors by enforcing expressing those molecules highly expressed in HSC but absent in downstream progenies. Ectopic expression of Sox17 can confer self-renewal potential on adult hematopoietic progenitors. However, this approach eventually led to leukemogenesis (He et al., 2011). Likewise, miR-125a is a non-coding RNA gene preferentially expressed in HSC rather than blood progenies (Guo et al., 2010). Ectopic expression of miR-125a in mouse hematopoietic progenitors induced long-term hematopoiesis, but the recipient mice suffered an MPN-like disease after secondary transplantation (Gerrits et al., 2012; Wojtowicz et al., 2016; Wojtowicz et al., 2014). Therefore, more extensive and innovative efforts are needed to develop safer approaches to convert blood progenitor cells into engraftable blood stem cells for ultimately therapeutic uses.

Hoxb5, a member of *HOX* gene family, is preferentially expressed in HSC, and uniquely marks the long-term HSC (Chen et al., 2016; Gulati et al., 2019). Our recent study showed that gain of function of Hoxb5 in pro-pre-B cells reprogrammed these cells into T lymphocytes *in vivo* (Zhang et al., 2018). Moreover, the latest research shows that exogeneous Hoxb5 expression confers protection against loss of self-renewal to Hoxb5-negative HSCs and can partially alter the cell fate of ST-HSCs to that of LT-HSCs (Sakamaki et al., 2021). Here, we further studied the potential role of Hoxb5 in MPP cell context, an intermediate progeny of HSC without self-renewal ability. Interestingly, conditional overexpression of Hoxb5 in MPP upon transplantation led to long-term hematopoiesis in serial transplanted mice. More importantly, Hoxb5 resulted in a *de novo* cell type defined as Mac1^+^CD48^+^SK, which contributed to the sustainable long-term hematopoiesis in serial transplant recipients. By single cell RNA-seq analysis, Mac1^+^CD48^+^SK cells demonstrated an activated machinery of DNA replication and cell division, resembling the characteristics of FL HSC, which associated with their acquired self-renewal feature. This study reveals *de novo* evidence that Hoxb5 can efficiently reprogram blood progenitors into engraftable blood stem cells, thereby offering a new strategy to expand the engraftable cell source for bone marrow transplantation

## Results

### Enforced expression of Hoxb5 in MPP leads to long-term hematopoiesis in transplantation setting

To evaluate the potential role of Hoxb5 in MPP, we chose *Hoxb5*^*LSL/+*^ Mx1-cre mouse model(Zhang et al., 2018) to conditionally express Hoxb5 in MPP. Upon transplantation into recipient animals, ectopic Hoxb5 expression can be turned on by injection with polyinosinic-polycytidylic acid (pIpC) and the GFP signal reports the expression of the Hoxb5 at single cell resolution. We sorted the conventional MPP (CD45.2^+^GFP^-^Lin (CD2, CD3, CD4, CD8, Mac-1, Gr1, B220, Ter119)^-^CD48^-^Sca1^+^c-kit^+^CD150^-^CD135^+^)(Adolfsson et al., 2001; Kiel et al., 2005) from the Sca1^+^ enriched bone marrow cells of *Hoxb5*^*LSL/+*^ Mx1-cre mouse or *Hoxb5*^*LSL/+*^ mouse. Four hundred sorted MPP along with 0.25 million Sca1^-^ helper cells (CD45.1^+^) were retro-orbitally transplanted into irradiated individual recipients (CD45.1^+^ C57BL/6 background) (***Figure 1A-B***). The recipients were injected intraperitoneally with pIpC (250 ug/mouse) every other day for six times starting at day5 before transplantation. We assessed the reconstitution ability of the donor derived cells by analyzing the peripheral blood (PB) chimeras every four weeks until 20^th^ week post-transplantation. (***Figure 1A***). Amazingly, in the primary recipients transplanted with the MPP of the *Hoxb5*^*LSL/+*^ Mx1-cre mouse, the ratio of the donor-derived cells (CD45.2^+^GFP^+^) continuously increased and the minimum ratio was up to 62% at the 20^th^ week post-transplantation, whereas the control recipients transplanted with the *Hoxb5*^*LSL/+*^ MPP shows a significantly low reconstitution ability, and the maximum donor-derived cells (CD45.2^+^) ratio was 11.4% at the 20^th^ week post-transplantation (***Figure 1C***). Furthermore, the contributions of donor-derived cells in spleen (SP) and bone marrow (BM) tissues of the Hoxb5-expressing MPP recipients were significantly more than the control group (*p* < 0.001) (***Figure supplement 1A***). In addition, multiple blood lineages including T cells (CD3^+^), B cells (CD19^+^), and Myeloid (Mac-1^+^or Gr1^+^) in the PB, SP, and BM were also detected at 20^th^ week post-transplantation in the primary recipients (***Figure 1D***). These results demonstrate that enforced expression of Hoxb5 in MPP leads to long-term hematopoiesis.

**Figure 1.**
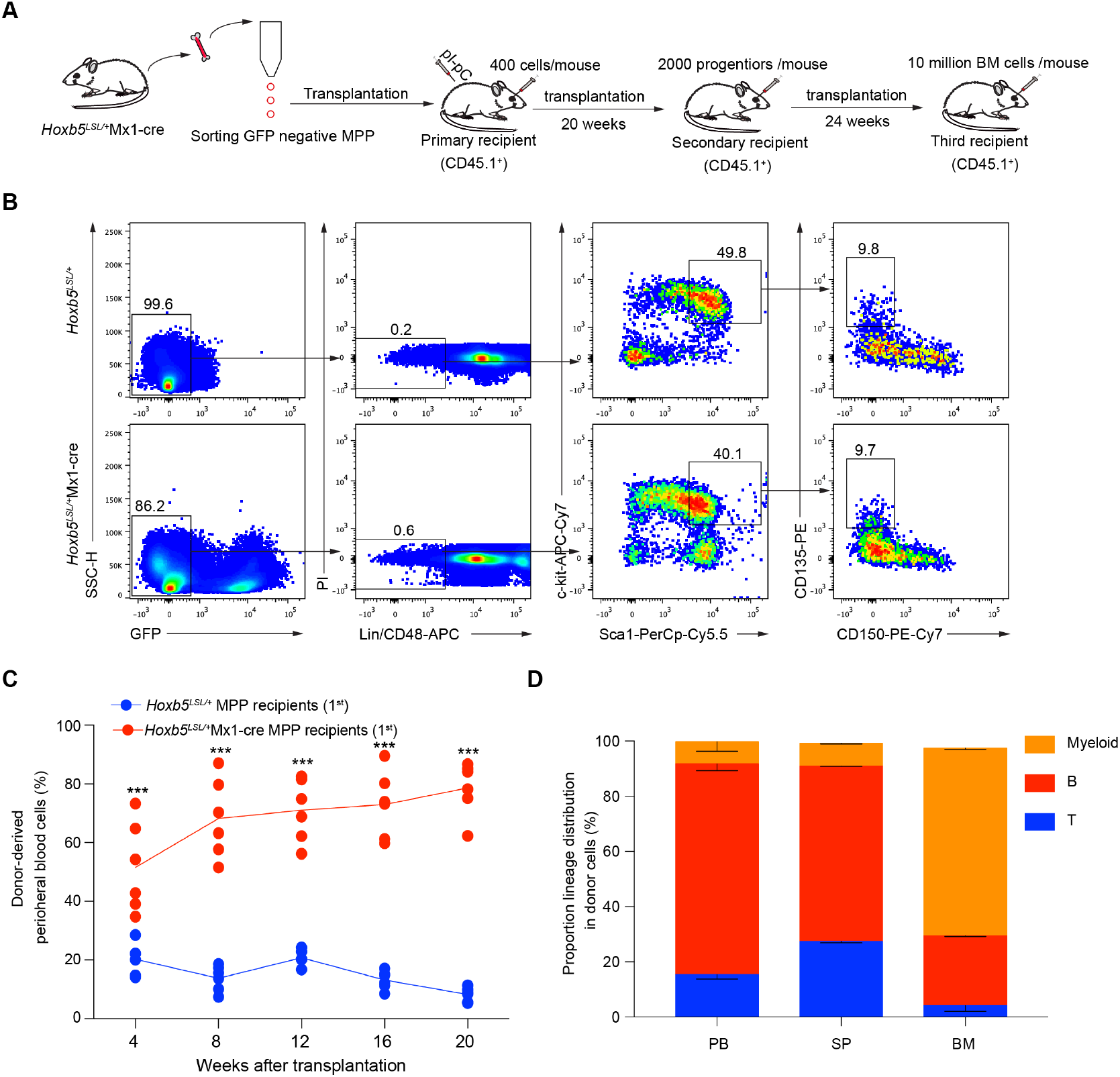
Overexpression of Hoxb5 impowers long-term reconstitution capacity on MPP. **(A)** The scheme for MPP transplantation. **(B)** Gating strategy for sorting the MPP. MPP were defined as CD45.2^+^GFP^-^Lin (CD2^-^CD3^-^CD4^-^CD8^-^Mac-1^-^Gr1^-^B220^-^Ter119^-^) CD48^-^Sca1^+^c-kit^+^CD150^-^CD135^+^ from either the *Hoxb5*^*LSL/+*^Mx1-cre mice or *Hoxb5*^*LSL/+*^ control mice (8 weeks old). The sorted MPP were transplanted into lethally irradiated (9.0 Gy) C57BL/6 mouse (CD45.1^+^, 400 cells/mouse) with CD45.1^+^Sca1^-^ helper cells (0.25million/mouse). Recipients were injected with pIpC (ip, 250 ug/mouse) every other day for six times starting from day 5 before transplantation. **(C)** Contribution curves of the donor-derived cells in peripheral blood cells (PB) of the primary recipients. The donor cells were defined as CD45.2^+^GFP^+^ (*Hoxb5*^*LSL/+*^Mx1-cre MPP recipients) or CD45.2^+^ (*Hoxb5*^*LSL/+*^ MPP recipients). The PB of the recipients transplanted with *Hoxb5*^*LSL/+*^Mx1-cre MPP (n = 6, as indicated by the red dot) or Hoxb5^LSL/+^ MPP (n = 6, as indicated by the blue dot) were analyzed every four weeks until the 20^th^ weeks after transplantation. mean ± SD, ***p< 0.001. Independent Samples *t* test. **(D)** Lineages distribution of the recipients (n = 3) at 20^th^ week after *Hoxb5*^*LSL/+*^Mx1-cre MPP transplantation. Columns shown are percentages of donor-derived T cells (CD3^+^), B cells (CD19^+^) and myeloid cells (Mac-1^+^ or Gr1^+^) in PB, spleen (SP), and bone marrow (BM).

### Hoxb5 results in the occurrence of a *de novo* Mac-1^+^CD48^+^ SK cell type associated with the long-term engraftable feature

To investigate the cellular mechanism, we analyzed the blood progenitor cells in the primary recipients at the 20^th^ week post-transplantation. We discovered a *de novo* donor-derived Sca1^+^c-kit^+^population cells, which simultaneously expressed Mac-1 and CD48 surface markers. Certainly, this cell type is not identified in natural blood cells in the absence of Hoxb5 expression (***Figure 2A***). Consistent with previous reports, natural MPP transplantation cannot sustainably give rise to LSK cells in the bone marrow of recipient mice (***Figure supplement 1B***). To further test whether the Mac-1^+^CD48^+^ SK cells are responsible for the long-term repopulating feature in Hoxb5 expressing MPP, we sorted the GFP^+^Mac-1^+^CD48^+^ SK cells and transplanted them into secondary recipient mice (CD45.1^+^ C57BL/6 background, 2000 cells/mouse) with Sca1^-^ helper cells (CD45.1^+^ 0.25 million/mouse). As expected, these Mac-1^+^CD48^+^ SK cells successfully reconstituted multi-lineage hematopoiesis in secondary recipients, as demonstrated by stable increases of donor-derived cells (GFP^+^) in the PB after transplantation (***Figure 2B***). Of note, the donor chimeras achieved as high as 94.6% at the 20^th^ week after transplantation and lineages of T, B and myeloid cells can be detected at the 4^th^, 12^th^ and 20^th^ post-transplantation (***Figure 2B-C***). Moreover, the donor-derived T, B and myeloid lineages in the PB, SP and BM also exhibited patterns resembling natural hematopoiesis at the 24^th^ week post-transplantation (***Figure 2D***). More importantly, the donor-derived Mac-1^+^CD48^+^ SK cells can still be detected in the BM of the secondary recipients (***Figure 2E***). These results indicate that the *de novo* Mac-1^+^CD48^+^ SK cell type is engraftable in the secondary recipients.

**Figure 2.**
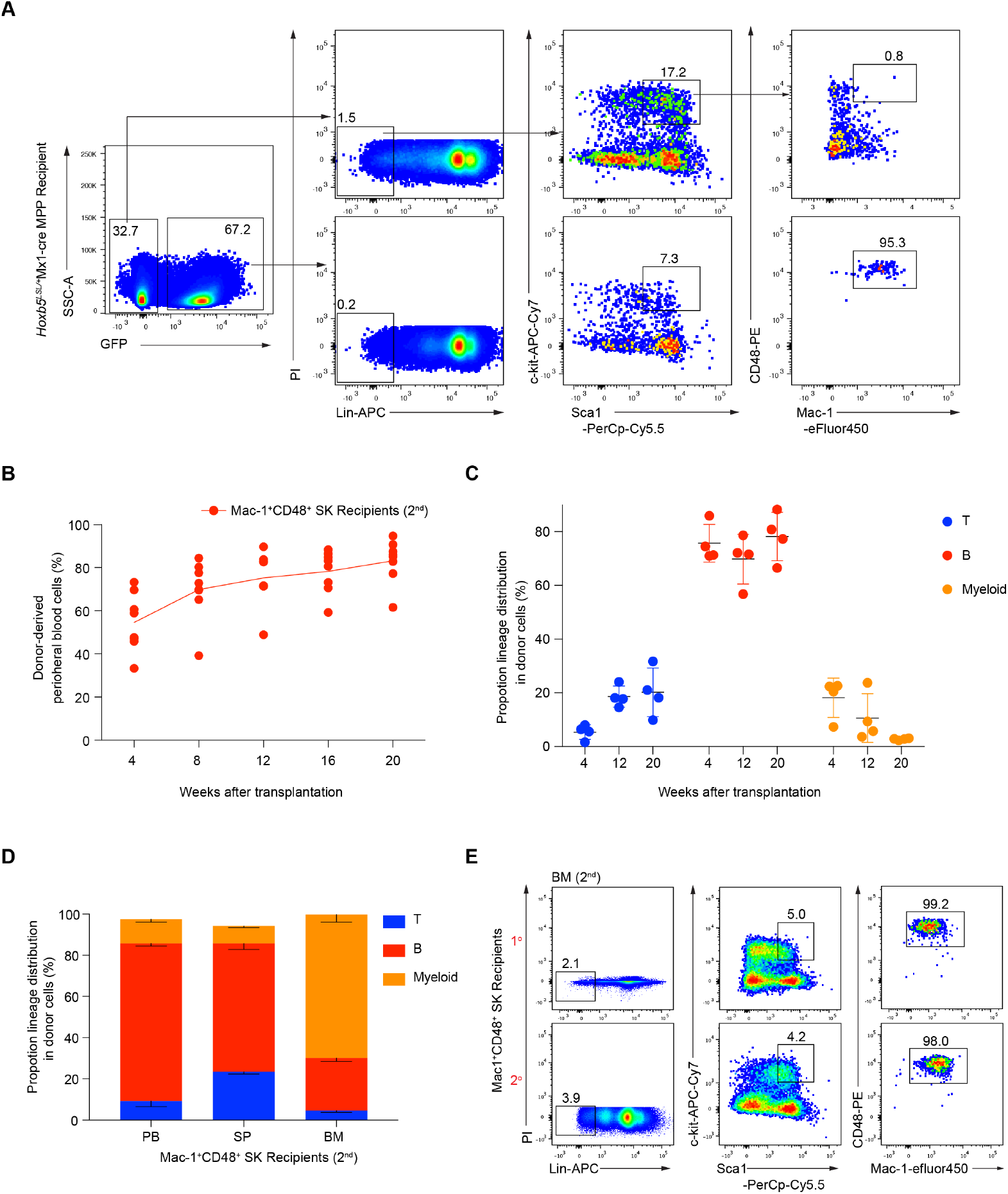
A *de novo* Mac-1^+^CD48^+^ SK cell population reconstitute hematopoiesis in secondary recipients. **(A)** FACS analysis of the donor-derived BM progenitors in the primary recipients 20 weeks post-transplantation. Antibodies of Lineages (CD2^-^CD3^-^CD4^-^CD8^-^Mac-1^-^Gr1^-^B220^-^Ter119^-^) (Lin), Sca1, c-kit, Mac-1 and CD48 were stained the BM of *Hoxb5*^*LSL/+*^Mx1-cre MPP recipients. Mac-1^+^CD48^+^ SK were defined as CD45.2^+^GFP^+^Lin^-^Sca1^+^c-kit^+^CD48^+^Mac-1^+^ were sorted from the primary recipients 20 weeks post-transplantation, and retro-orbitally transplanted into secondary recipients (9.0 Gy, 2000 cells/mouse). **(B)** Chimeras curves of the donor cells to the peripheral blood (PB) cells of the secondary recipients (n = 7). For the secondary transplantation, Mac-1^+^CD48^+^ SK cells were retro-orbitally injected into the lethally irradiated recipients (9.0 Gy, 2000 cells/mouse). The donor-derived cells (CD45.2^+^GFP^+^) in the PB were analyzed every four weeks post-transplantation. **(C)** Lineages distribution in PB of the secondary recipients (n = 4) at the week-4^th^, 12^th^ and 20^th^ post-transplantation. Proportions of the CD3^+^ (T), CD19^+^ (B), Mac1^+^ (Myeloid) in donor-derived cells were analyzed. **(D)** Lineages distribution of the recipients (n = 3) at 24^th^ week after Mac-1^+^CD48^+^ SK transplantation. Columns shown are percentages of donor-derived T cells (CD3^+^), B cells (CD19^+^) and myeloid cells (Mac-1^+^ or Gr1^+^) in PB, spleen (SP), and bone marrow (BM). **(E)** Immuno-phenotypes of the donor-derived Mac-1^+^CD48^+^ SK in the bone marrow (BM) of the secondary recipients. Two representative mice were shown.

To assess the long-term hematopoiesis capacity of the Mac-1^+^CD48^+^ SK cells. we did the third transplantation using total BM cells of the Mac-1^+^CD48^+^ SK recipients (Secondary recipients). Recipients (CD45.1^+^ C57BL/6 background) accepted the lethally irradiation first (9 Gy), and then were retro-orbitally injected with the total BM cells (10 million/mouse, n = 6) from the secondary recipients. The contribution of CD45.2^+^GFP^+^ donor cells to the peripheral blood was assessed at week-8^th^, 16^th^, 20^th^, 26^th^ and 32^th^ after transplantation. All of the recipients were reconstituted with the CD45.2^+^GFP^+^ cells with the ratio of 48.7%-74.2% 8 weeks post-transplantation, and the average ratio was still 49% after transplantation for 32 weeks (***Figure 3A***). Moreover, the donor-derived cells ratio even has no significant difference (*p* = 0.075) even at the week 32^th^ compared with the week 8^th^ post-transplantation (***Figure 3B***). Meanwhile, the donor-derived cells also showed multi-lineage distributions in PB after 8 weeks and 20 weeks later post-transplantation (***Figure 3C***). Furthermore, The average ratio of T cells (CD3^+^) was 7.1% (week 8^th^, n = 6), 17.4% (week 20^th^, n = 6) and 16.1% (week 32^th^, n = 6), B cells (CD19^+^) was 83.2% (week 8^th^, n = 6), 76.5% (week 20^th^, n = 6) and 82.8% (week 32^th^, n = 6), and Myeloid cells (Mac1^+^ or Gr1^+^) was 11.1% (week 8^th^, n = 6), 10.6% (week 20^th^, n = 6) and 7.7% (week 32^th^, n = 6) post-transplantation in PB respectively (***Figure 3D***). Collectively, these results indicate that the Mac-1^+^CD48^+^ SK cells can sustain the long-term hematopoiesis in serial transplantation.

**Figure 3.**
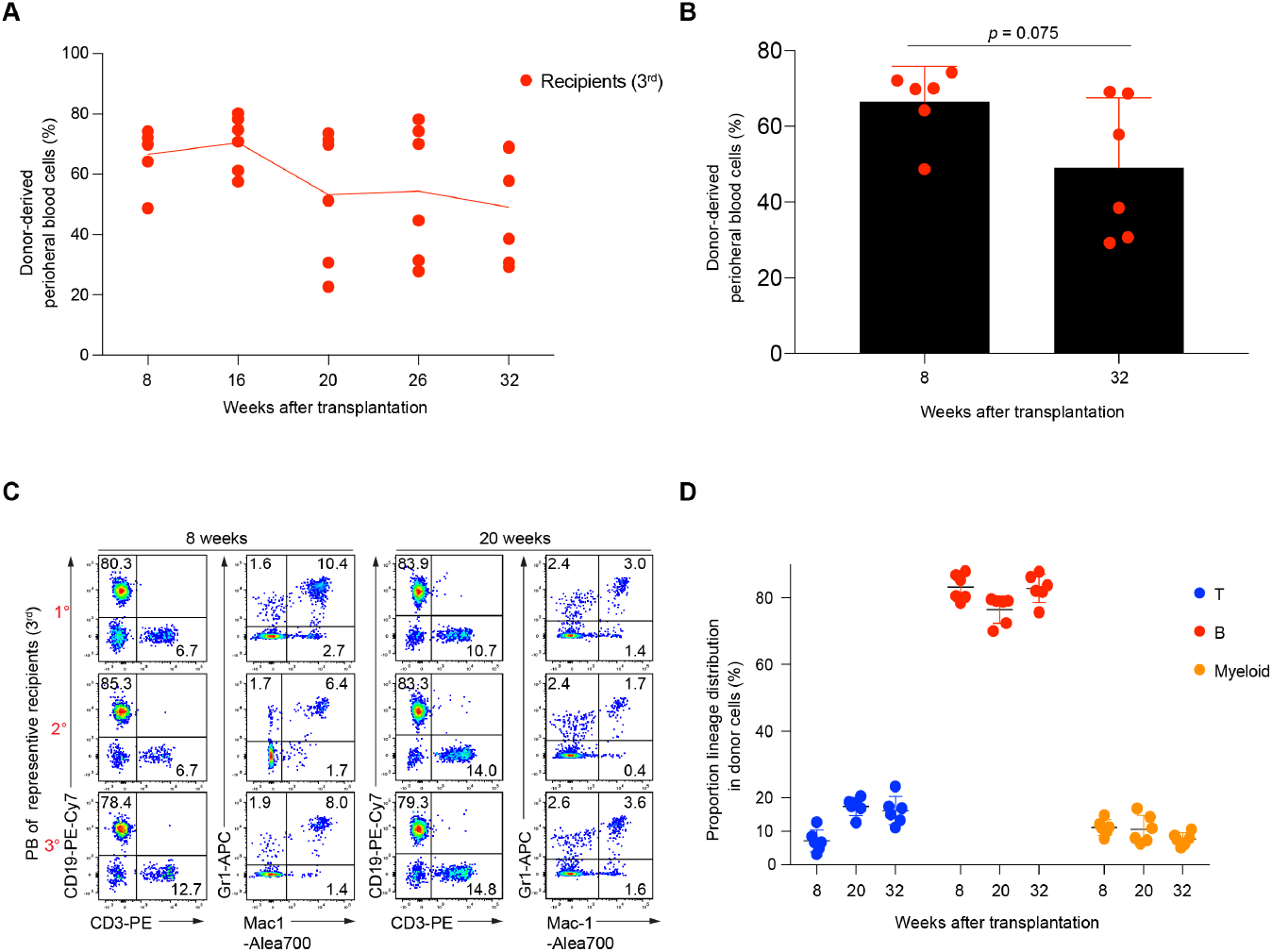
Mac-1^+^CD48^+^ SK cells still maintained the repopulation capacity in the third transplantation recipients. **(A)** Chimeras curves of the donor cells to the PB of the secondary recipients (n = 6). Donor-derived cells (CD45.2^+^GFP^+^) in the PB were analyzed at 8^th^, 16^th^, 20^th^, 26^th^ and 32^th^ week post-transplantation. For the third transplantation, recipients (CD45.1^+^ C57BL/6) were lethally irradiated (9 Gy) and then were retro-orbitally injected with the nucleated BM cells (10 million/mouse) isolated from the secondary recipients. **(B)** Comparation of the donor derived cells (CD45.2^+^GFP^+^) ratio at 8 weeks and 32 weeks post-transplantation. Independent Samples *t* test. **(C)** Representative FACS analysis (n = 3) of the PB from the third transplantation recipients (3^rd^) after transplanting with the total BM cells of the secondary recipients 8 weeks and 20 weeks later. **(D)** Lineages distribution in PB of the third recipients (n = 6) at the week-8^th^, 20^th^ and 32^th^ post-transplantation. Proportions of the CD3^+^ (T), CD19^+^ (B), Mac1^+^or Gr1^+^ (Myeloid) in donor-derived cells were analyzed.

### Mac-1^+^CD48^+^ SK cells showed proliferating signatures of DNA replication and cycling, resembling fetal liver HSC cells

To investigate the underlying molecular mechanism, we characterized the Mac-1^+^CD48^+^ SK cells (n = 47) at transcriptome level by single cell RNA-seq analysis. Meanwhile, we also performed the single cell RNA-seq of the BM HSC (n = 36, Hoxb5^LSL/+^ mice, 8 weeks old) and WT-MPP (n = 42, Hoxb5^LSL/+^ mice, 8 weeks old). Certainly, we also sorted the single cells of FL HSC (n = 56, Hoxb5^LSL/+^ mice, Day14.5 embryo), and performed the RNA-seq (***Figure supplement 1C***). To dissect the transcriptome signature between Mac-1^+^CD48^+^ SK cells and the other three cell types (BM HSC, FL HSC, WT-MPP), we first found out the differentially expressed genes (adjusted *P* value <0.05) between FL HSC and WT-MPP. The up- and down-regulated differential expressed genes for FL HSC versus WT-MPP were respectively used as gene set for Gene set-enrichment analysis (GSEA) between Mac-1^+^CD48^+^ SK cells and WT-MPP. (***Figure 4A-B, File supplement 1-2***). The results showed that the up-regulated genes were enriched in the Mac-1^+^CD48^+^ SK cells (***Figure 4A***). Meanwhile, we also calculated out the up- and down-regulated differential expressed genes (adjusted P value <0.05) for FL HSC versus BM HSC. The GSEA between Mac-1^+^CD48^+^ SK cells and WT-MPP suggested that these up-regulated genes were also enriched in the Mac-1^+^CD48^+^ SK cells (***Figure 4C-D, File supplementary 3-4***). Moreover, We combined the leading edge genes from Fig.4a and Fig.4c, and performed the heatmap analysis. The result showed that the expression level of these genes in Mac-1^+^CD48^+^ SK cells is equivalent to these in FL HSC cells (***Figure 4E***). We further performed Gene-ontology (GO) analysis using these leading edge genes, and observed that they are abundantly involved in cell proliferation process, especially the processes of DNA replication, chromosome segregation (***Figure 4F***). Moreover, besides the higher expression of Hoxb5 both in Mac-1^+^CD48^+^ SK cells and FL HSC compared with BM HSC and WT-MPP, several genes were also up-regulated, which not only regulating the cell cycle, but also have a vital role in the regulation of hematopoiesis, including Birc5 (Gurbuxani et al., 2005), *Gmnn* (Yasunaga et al., 2016), Cdt1 (Yasunaga et al., 2016), *Cdc45* (Flach et al., 2014), and *Gins1* (Ueno et al., 2009) (***Figure 4G***). Furthermore, we also find out the genes of Cdk6 (Scheicher et al., 2015), Satb1 (Will et al., 2013; Yasui et al., 2002), Runx3 (de Bruijn and Dzierzak, 2017) and *Mybl2* (Baker et al., 2014; Bayley et al., 2018), which were only up-regulated in Mac-1^+^CD48^+^ SK cells, and have an important role in HSC homeostasis or development. Thus, these results indicate that Hoxb5 expression empowers self-renewal capacity on Mac-1^+^CD48^+^ SK cells by activating cell cycle and DNA replication machinery, resembling the FL HSC cells.

**Figure 4.**
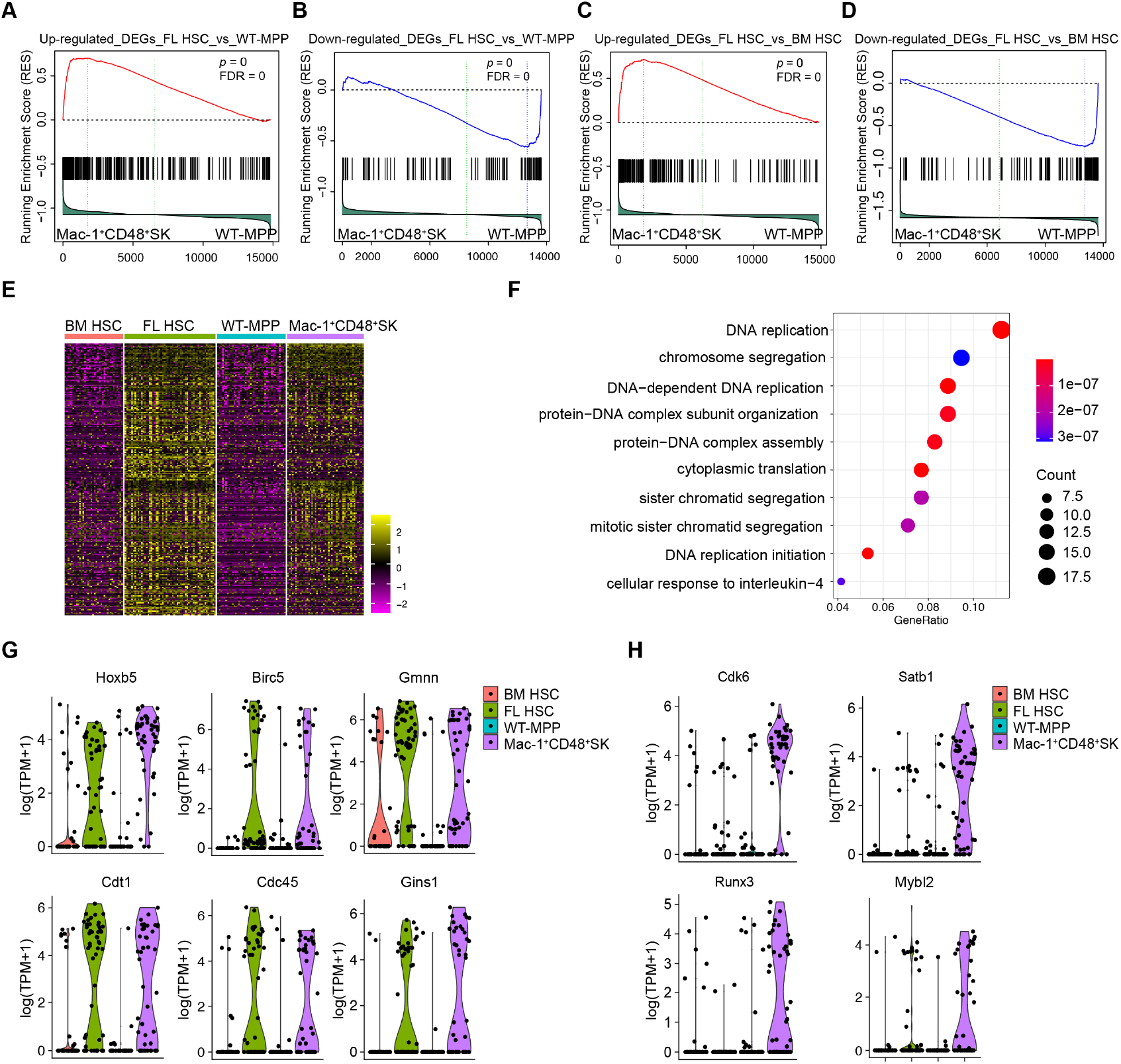
Rapid proliferation features of Mac-1^+^CD48^+^ SK cells at single cell resolution. **(A)** Gene set–enrichment analysis of WT-MPP (n = 42) and Mac-1^+^CD48^+^ SK (n = 47). The gene set used to analysis were from the up-regulated genes in FL HSC (n = 56) versus WT-MPP (n = 42) (adjusted P value <0.05). **(B)** Gene set–enrichment analysis of WT-MPP (n = 42) and Mac-1^+^CD48^+^ SK (n = 47). The gene set used to analysis were from the down-regulated genes in FL HSC (n = 56) versus WT-MPP (n = 42) (adjusted P value <0.05). **(C)** Gene set– enrichment analysis of WT-MPP (n = 42) and Mac-1^+^CD48^+^ SK (n = 47). The gene set used to analysis were from the up-regulated genes in FL HSC (n = 56) versus BM HSC(n = 36) (adjusted P value <0.05). **(D)** Gene set–enrichment analysis of WT-MPP (n = 42) and Mac-1^+^CD48^+^ SK (n = 47). The gene set used to analysis were from the down-regulated genes in FL HSC (n = 56) versus BM HSC(n = 36) (adjusted P value <0.05). **(E)** Heatmap analysis of the BM HSC, FL HSC, WT-MPP and Mac-1^+^CD48^+^ SK. Genes used to analysis were from the leading edge genes in **(A)** and **(C). (F)** Gene ontology (GO) enrichment analysis of genes from the leading edge genes in **(A)** and **(C)**. Each symbol represents a GO term (noted in plot); color indicates adjusted P value (Padj (significance of the GO term)), and symbol size is proportional to the number of genes. **(G)** Violin plots show the expression profile of selected factors (Hoxb5, Birc5, Gmnn, Cdt1, Cdc48 and Gins1) both in Mac-1^+^CD48^+^ SK and FL HSC related to DNA replication, cell division and hematopoiesis at single cell resolution. **(H)** Violin plots show the expression profile of selected factors (Cdk6, Satb1, Runx3 and Mybl2) preferentially expressed in Mac-1^+^CD48^+^ SK related to hematopoiesis at single cell resolution.

## Discussion

In this study, we explored the role of Hoxb5 in MPP cell context. Hoxb5 expression leads to long-term hematopoiesis of MPP in serial transplantation settings by generating the Mac-1^+^CD48^+^ SK cells, a *de novo* cell type that naturally does not exist. At the transcriptome level, the Mac-1^+^CD48^+^ SK cells showed molecular signatures of cell division and proliferation resembling the FL HSC, which correlating their acquired self-renewal feature.

Stemness feature is the only functional difference between HSC and their progeny MPP. However, the stemness-losing mechanism along the differentiation path from HSC to MPP is unknown. The genes shut down or down-regulated in MPP, such as *Hoxb5*, might be accountable for the loss of stemness from HSC to MPP. Moreover, this maybe indicated by the latest research, which reported that exogeneous Hoxb5 expression confers protection against loss of self-renewal to Hoxb5-negative HSC and can partially alter the cell fate of ST-HSC to that of LT-HSC (Sakamaki et al., 2021). Here, despite over-expressing Hoxb5 in MPP generated no phenotypic HSC-like cells, the *de novo* Mac-1^+^CD48^+^ SK cells can substitute natural HSC for long-term engraftment. Interestingly, FL HSC shared two features with Hoxb5-expressing Mac-1^+^CD48^+^ SK cells, one is undergoing rapid proliferation, and the other is expressing Mac-1 marker (Kim et al., 2006; Morrison et al., 1995). However, natural adult HSC lose the expression of Mac-1 (Morrison and Weissman, 1994), which is consistent with their predominant dormancy under homeostasis. Therefore, the expression of Mac-1 is phenotypically associated with the fast expanding features of FL HSC and Hoxb5-expressing Mac-1^+^CD48^+^ SK cells. Seemingly, the enforced expression of Hoxb5 in MPP activates a cell division machinery (Dalton, 2015; Gao and Liu, 2019) without compromising their multi-lineage differentiation potential just as FL HSC.

Reportedly, ectopic expression of either Sox17 or miR-125a in MPP can confer a self-renewal ability, but eventually resulted in hematologic malignancies (Chhabra and Mikkola, 2011; Hu and Shivdasani, 2005; Krivtsov et al., 2006). MiR-125a-induced MPN displayed a complex manner of oncogene-dependency (Guo et al., 2012). Interestingly, no hematologic malignancies were found in the recipients transplanted with the Hoxb5-expressing MPP. Even the expression of Hoxb5 in total BM of the *Hoxb5*^*LSL/+*^ vav-cre mouse showed a normal hematopoiesis (Zhang et al., 2018). Thus, the self-renewal feature activated by Hoxb5 might be insulated from oncogenesis.

We also tested the engraftment potential of HOXB5-expressing human MPP in immunodeficient animals. Unfortunately, the HOXB5-expressing human MPP failed to recapitulate the long-term engraftment phenotype of Hoxb5-expressing murine MPP (data not shown). One possible reason is that the function of HOXB5 is not conservative between human and mouse species. However, we cannot exclude another possibility that HOXB5-overexpressing human MPP need a humanized bone marrow micro-environment for HOXB5-reprgramming, which is not available in current immunodeficient animal models.

In conclusion, our study reveals a rare role of Hoxb5 in empowering self-renewal capacity on MPP, which provides insights into converting blood progenitors into alternative engraftable cell source.

### Materials and methods Mice

Animals were housed in the animal facility of the Guangzhou Institutes of Biomedicine and Health (GIBH). *Hoxb5*^*LSL/+*^ mice were described as previous reported (Zhang et al., 2018). CD45.1, Mx1-cre and Vav-cre strains were purchased from the Jackson laboratory. All the mouse lines were maintained on a pure C57BL/6 genetic background. All experiments were conducted in accordance with experimental protocols approved by the Animal Ethics Committee of GIBH.

### Flow cytometry

Antibodies to CD2 (RM2-5), CD3 (145-2C11), CD4 (RM4-5), CD8a (53-6.7), Gr1 (RB6-8C5), Mac-1 (M1/70), Ter119 (TER-119), B220 (6B2), c-kit (2B8), Sca-1 (E13-161.7), CD135 (A2F10), CD150 (TC15-12F12.2), CD19 (eBio1D3), CD48 (HM48-1) ki-67 (16A8), Fcγ RII/III (2.4G2), CD127 (SB/199), CD45.2 (104) CD45.1(A20) were purchased from eBioscience or BioLegend. DAPI, 7-AAD and PI were used to stain dead cells. Flow cytometry was performed on an LSR Fortessa (BD Biosciences) and data were processed by FlowJo software (Version: 10.4.0, Tree Star).

### Cell sorting

Cells used for sorting were first incubated with the biotin-conjugated antibody to Sca1 (anti-Sca1 biotin) and then enriched using Anti-Biotin MicroBeads by AutoMACS Pro (Miltenyi Biotec). The enriched cells, stained with the antibodies, were sorted by BD FACSAria III.

### Transplantation

All recipients (CD45.1^+^, C57BL/6) were lethally irradiated (9 Gy, RS2000, Rad Source) at least 4 hours before transplantation. MPP (400 cells/mouse) from Hoxb5^LSL/+^ mouse or Hoxb5^LSL/+^ Mx1-cre mouse for primary transplantation, and donor-derived CD48^+^Mac-1^+^ SK (2000 cells/mouse) from the primary recipients for secondary transplantation were retro-orbitally transplanted into the recipients with the Sca1^-^ helper cells (CD45.1^+^ 0.25 million/mouse). For third transplantation, total BM cells (10 million/mouse) of the secondary recipients were used as the donor cells. To induce Hoxb5 expression, the primary recipients were intraperitoneally injected with polyinosinic-polycytidylic acid (pIpC) (250 ug/mouse) every other day for six times starting from the day5 before transplantation. Recipients were fed with the water added with trimethoprim-sulfamethoxazole for one month after irradiation.

### RNA-seq and data analysis

cDNA of the single cell from adult wide type HSC (BM HSC, Hoxb5^LSL/+^ mice, 8weeks old), Fetal liver HSC (FL HSC, Hoxb5^LSL/+^ mice, Day14.5, defined as CD45.2^+^Lin^-^Sca1^+^c-kit^+^Mac1^+^CD150^+^), Wide type MPP (WT-MPP, Hoxb5^LSL/+^ mice, 8 weeks old) and donor-derived CD48^+^Mac-1^+^ SK cells (Primary recipients, week 8^th^ post-transplantation) were generated and amplified using Discover-sc WTA Kit V2 (Vazyme). B2m and Gapdh were used to assess the quality of the amplified cDNA by qPCR analysis. The qualified samples were used to prepare the sequencing library by the TruePrep DNA Library Prep Kit V2 (Vazyme), and the qualified libraries were sequenced by illumina sequencer NextSeq 500. Raw data (fastq files) were generated using bcl2fastq software (version 2.16.0.10) and were uploaded to the Gene Expression Omnibus public database (GSE NO.183800). HISAT2 (version 2.1.0) were used to align the raw data, and the StringTie (version 1.3.4) were used to estimate the expression level in the transcripts per million (TPM) as previously reported(Pertea et al., 2016; Pertea et al., 2015). The DESeq2 was used for differential expression analysis, and the related Heatmaps were potted using pheatmap (version 1.0.8). Principle Component Analysis (PCA) were used by prcomp function of stats (R package, version 3.4.4) and PCA plots and violin plots were plotted using ggplot2 (R package, version 2.2.1). Gene set-enrichment analysis (GSEA) and gene-ontology (GO)-enrichment analysis (clusterProfiler package) were performed as described(Subramanian et al., 2005; Yu et al., 2012).

## Acknowledgments

This work was supported by grants from the grants from the National Natural Science Foundation of China (Grant No. 82100127, 81925002, 31900816).

## Author Contributions

J.W., F.H., and Q.Z. designed the project. F.H., Q.Z. and D.H conducted all the experiments and data analysis. M.Z. performed the RNA-seq experiments, and Q.W. analyzed the RNA-seq data. Q. Z. performed part of the mouse genotype experiments. K.W., Q.W., and J.X. performed the irradiation experiments, L. L., C.X., and T.W. participated in the Human cord blood MPP experiments. X.L. and Y.G. setup the flow cytometry. Y.G. ordered the experiment reagents. H.C., F.D., F.H. and J.W discussed the data. F.H. and J.W. wrote the manuscript and approved it

## Conflict of Interest

The authors declare there’s no competing financial interests in relation to the work described

---

**Figure supplement 1.**
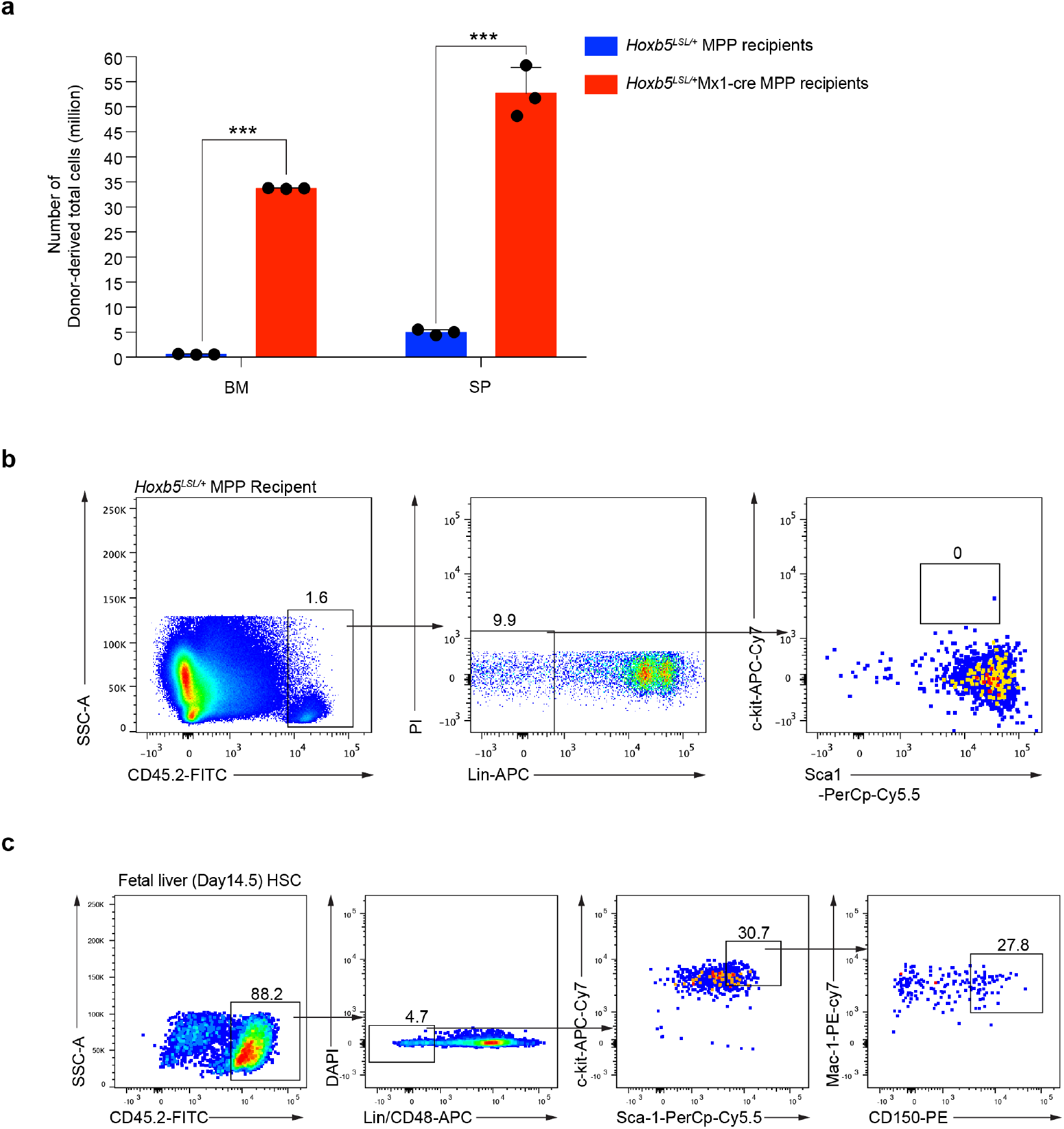
**(A)** Statistic analysis of the absolute cell counts in the bone marrow (BM) and spleens (SP) of the donor-derived cells from the *Hoxb5*^*LSL/+*^ MPP recipients or *Hoxb5*^*LSL/+*^Mx1-cre MPP recipients. mean ± SD, \*\*\**p*<0.001. Independent Samples *t* test. **(B)** FACS analysis of the donor-derived LSK cells from *Hoxb5*^*LSL/+*^ MPP recipients 20 weeks post-transplantation. Antibodies of Lineages (CD2^-^CD3^-^CD4^-^CD8^-^Mac-1^-^Gr1^-^B220^-^Ter119^-^) (Lin), Sca1, c-kit, Mac-1 and CD48 were stained the BM of the recipients. **(C)** Sorting strategy of the FL HSC. The fetal livers were dissected from embryos (day14.5), and the antibody cocktail stained nucleated cells were used to sorting FL HSC (DAPI^-^ CD45.2^+^Lin(CD2 CD3 CD4 CD8 Gr1 Ter119 B220 IgM CD48)^-^Sca-1^+^ckit^+^Mac1^+^CD150^+^).

## Notes

### Competing Interest Statement

The authors have declared no competing interest.

